# Cellular context defines brain-specific functions of genes associated with neurodevelopmental disorders

**DOI:** 10.64898/2026.10.26.761783

**Authors:** Milena Djokic, Sivarajan Karunanithi, Mareen Welzel, Fridolin Kielisch, Katja Luck

## Abstract

Differential gene expression rarely explains why certain genes, e.g. associated with tissue-specific manifestations of human disease, exert tissue-specific functions. To understand how gene expression and molecular context act in concert to drive tissue-specific gene function, we focus on Neurodevelopmental disorders (NDDs), for which underlying disease mechanisms are still poorly understood. We developed a computational framework that harmonizes and integrates spatiotemporal transcriptome datasets enabling us to systematically quantify preferential gene expression across tissues and developmental stages. Applying this pipeline to approximately 1,000 NDD-associated genes, we find that only about 10% show brain-preferential expression, with distinct gene clusters instead exhibiting significant upregulation in non-brain tissues. Investigation of these clusters reveals regulatory mechanisms that govern mitochondrial function to meet neurons’ high energetic demands, which are mechanistically distinct from those employed by other energy-intensive tissues such as heart and skeletal muscle. Furthermore, by integrating tissue-agnostic protein-protein interaction data with our expression resource, we identify putative brain-specific interaction partners for 22 uniformly expressed NDD-associated genes. We experimentally validate these interactions for QRICH1 and demonstrate that disease-associated mutations disrupt them, providing a molecular hypothesis for how QRICH1 acquires brain-specific function and how its disruption might cause Ververi-Brady syndrome. Together, our findings highlight the importance of molecular context for brain-specific gene function in the context of neurodevelopmental disease. We provide a framework for dissecting molecular mechanisms of tissue-specific Mendelian-like disorders and for exploring how biological processes are adapted to tissue-specific needs.

## Introduction

Multicellular organisms consist of hundreds of different cell types and dozens of different tissues. Despite their striking phenotypic differences, these cells and tissues share largely identical genetic information. Understanding the molecular mechanisms that drive this functional diversity at the cell and tissue level is one of the most fascinating puzzles in biology. Two major concepts emerged^1^. First, various cellular processes can generate cell or tissue-specific gene products, which can then exert cellular context-specific functions. These are processes such as differential gene expression, alternative splicing, and posttranslational modifications as well as differential subcellular localization of gene products. Of course, for the synthesis of cellular context-specific gene products, cellular context-specific mechanisms need to be in place, such as specifically expressed transcription factors, splice factors, or kinases. Second, gene products might not be themselves context-specific but interact with cellular context-specific partner molecules either directly or indirectly, i.e. by being part of a protein complex with other context-specific subunits. Such context-specific interactions would then enable the execution of context-specific functions by the otherwise uniformly expressed gene product.

Contrary to common belief, various studies have revealed that tissue-specific expression is not a major mechanism by which gene products adopt tissue-specific functions. For instance, large scale tissue transcriptome studies reported less than 2,000 genes found to be expressed in only one tissue^2^. Other studies estimated that number to range between 15 and 30%^3^. Interestingly, 40% of genes with uniform expression across tissues are annotated with tissue-specific gene ontology terms supporting that other mechanisms than gene expression frequently drive the adoption of tissue-specific functions^4^. Previous studies have systematically investigated the possibility for proteins to adopt tissue-specific functions via interaction with tissue-specific partner proteins^3,5–7^. Via integration of cellular context-agnostic protein-protein interaction (PPI) data with gene expression data, it was shown that the fraction of the protein-protein interactome that is likely tissue-specific is substantially larger than the fraction of the genome expressed in a tissue-specific manner. This is because any interaction mediated by a tissue-specific protein is considered tissue specific, regardless of the expression of its partner^6,7^. While trivial, this insight underlines the importance of molecular context for the generation of cellular context-specific function.

Further insights into mechanisms that confer tissue-specific functions to genes have been gained from studying Mendelian-like disorders. These heritable diseases are commonly caused by a single pathogenic germline variant, which is, thus, present in every cell of the affected individuum. Yet, an analysis of 1,200 Mendelian disorders revealed that about two third exhibit very tissue-specific phenotypes^4^. While the corresponding disease-associated genes showed on average elevated expression in these affected tissues, 80% of these genes are also well expressed in other non-affected tissues. Previous studies have systematically interrogated PPI networks formed between disease-associated uniformly expressed proteins and tissue-specific partner proteins to find that disease-associated proteins exhibited a higher median number of tissue-specific PPIs in the affected tissue compared to unaffected tissues, although it remains unclear to which extent this observation is driven by a potential study bias^3^. Others have discovered potential disease-relevant tissue-specific PPIs and experimentally showed that these are disrupted by disease-associated mutations providing a clear molecular hypothesis for a mechanism that might drive disease^7^. However, these principles, while being broadly studied, have not been applied to advance our understanding of molecular mechanisms for selected diseases.

Neurodevelopmental disorders (NDDs) are a group of Mendelian-like diseases that are characterized by the disruption of brain developmental processes such as neurogenesis, neuronal differentiation, neuronal migration, and the formation and signaling of synapses^8^. These impairments manifest as deficits in cognitive, behavioural, and motor functions and result in disorders such as epilepsy, developmental delay, intellectual disability, and autism spectrum disorders, as well as numerous rare diseases. Although they are individually uncommon, about 3% of all children worldwide are diagnosed with an NDD^9^. NDDs predominantly affect brain function, yet, it remains unclear if loss-of-function of brain-specifically expressed genes is a major mechanism that drives NDDs or to which extent the molecular context of NDD-associated genes dictates their brain-specific functions.

Importantly, NDDs are likely caused by molecular perturbations in the brain during prenatal development requiring the analysis of pre- and postnatal gene expression across tissues to connect differential expression patterns of NDD genes with putative disease mechanisms. The Allen Brain Atlas^10^ is a unique resource that profiles gene expression across brain subregions as well as various pre- and postnatal developmental stages, however, because it is restricted to the brain, it cannot be used to assess differential expression of NDD genes across tissues. GTEx^2^, on the other hand, is unique in its breadth and depth for providing transcriptome data across adult tissues precluding though prenatal samples. Other studies profiled gene expression at various pre- and postnatal stages for a handful of tissues, yet consisted of small sample sizes that might hinder quantification of preferential expression^11^. Thus, systematic analysis of spatiotemporal gene expression patterns of NDD genes requires computational pipelines that can integrate highly heterogeneous gene expression datasets and quantify tissue preferential expression across those.

To overcome these challenges we developed a computational framework that re-processes large tissue transcriptome datasets in a tissue-aware manner and quantifies tissue-preferential expression across tissues and developmental stages. We used the resulting gene expression resource to assess tissue-preferential expression patterns of about 1,000 NDD-associated genes to find that only about 10% display brain-preferential expression. To our surprise, we obtained distinct clusters of NDD genes with significant upregulation in other tissues than the brain. Further investigation of these clusters and associated processes gives support for molecular mechanisms that regulate mitochondrial function in neurons to fulfil their high energy demands. We show that these mechanisms are distinct from how other energy-demanding tissues such as heart and skeletal muscle adapt mitochondria function. Furthermore, we identify brain-specific interaction partners for 22 uniformly expressed NDD-associated genes, experimentally confirm those for QRICH1, and probed their disruption by disease-associated mutations. This provides first hypotheses for molecular mechanisms by which QRICH1 adopts brain-specific functions and causes Ververi-Brady syndrome upon their disruption.

## Results

### Quantification of tissue-preferential gene expression

To explore the spatiotemporal expression patterns of NDD-linked genes across human tissues and developmental stages we searched for RNA-seq datasets that sampled gene expression across prenatal and postnatal developmental stages for many tissues including the brain. We selected the GTEx dataset^2^, which contains 17,382 RNA-seq samples across 52 tissues from different postnatal stages (Figure 1A, S1A). The number of samples and standardization of sample taking and measuring resulted in a highly robust resource to quantify preferential expression across human adult tissues. To quantify tissue-preferential expression of genes in prenatal stages, we chose a dataset from Cardoso-Moreira et al. (hereafter referred to as the Kaessmann dataset, Figure 1A)^11^. The Kaessmann dataset consists of 313 samples taken from 7 tissues in 13 pre- and 8 postnatal stages (Figure S1A). We reprocessed both RNA-seq datasets using the R package YARN^12^ to perform quantile normalization while preserving biological differences between tissues and developmental stages. Multidimensional scaling (MDS) analysis revealed high similarity between some tissues such as 13 brain subregions in GTEx that were consequently regrouped into the three larger brain regions cerebellum, basal ganglia, and others (Figure S1B), similar to previously published work^13^. Similarly, we found that the pre- and postnatal samples of each tissue profiled in the Kaessmann dataset clustered together (Figure 1B). Based on this observation, we combined these samples into one prenatal (PrenatK) and one postnatal (PostnatK) developmental stage, each containing four tissues.

**Figure 1.**
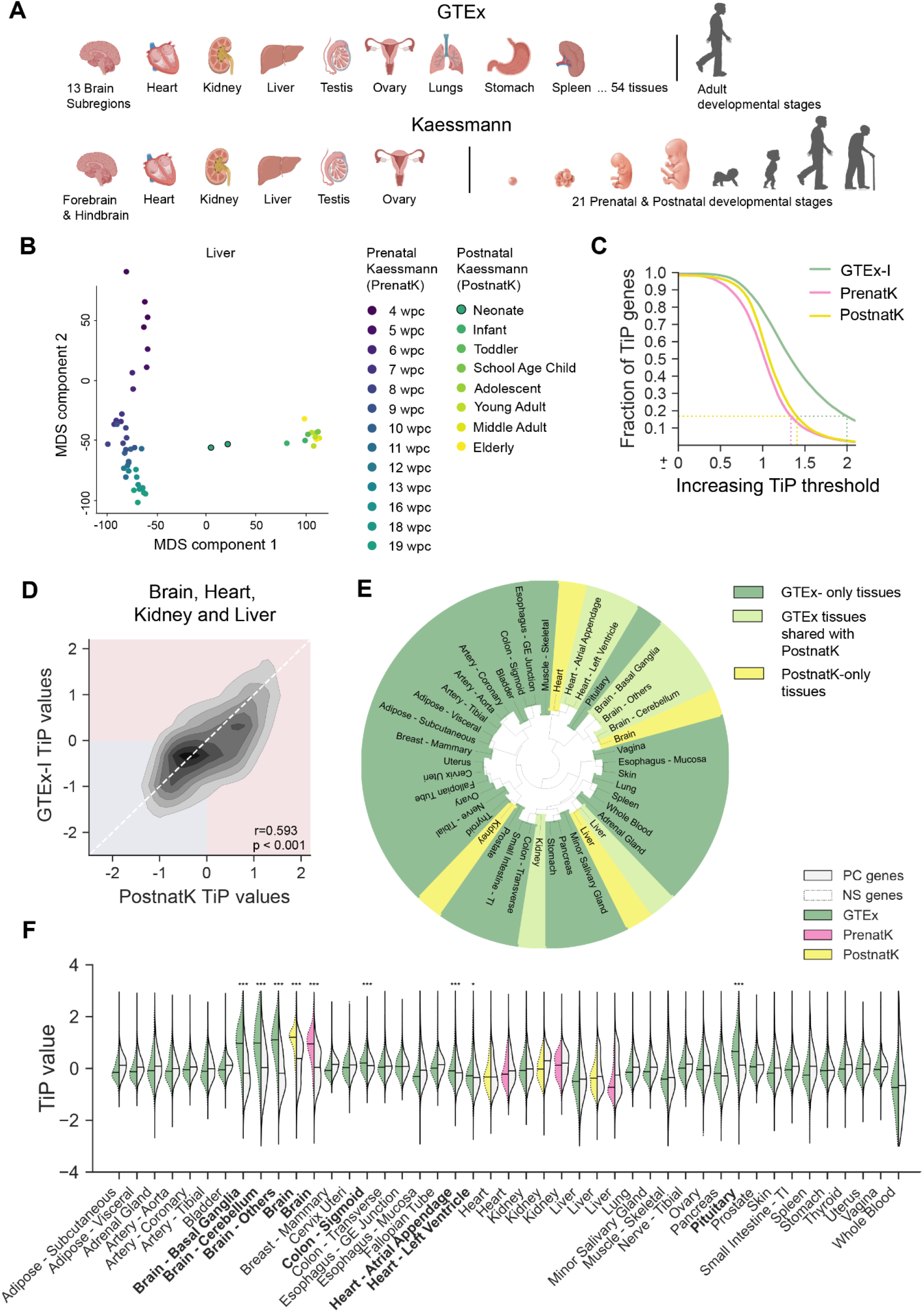
Processing of gene expression datasets and TiP value computation. **A.** Schematic illustration of the tissues and developmental stages of samples from the GTEx and Kaessmann resource. **B.** MDS plot of liver samples from the Kaessmann resource. **C.** Line plot showing the number of TiP genes for increasing TiP value cutoffs for different gene expression resources. Dashed lines indicate the cutoffs chosen for each dataset. **D.** Density plot showing the correlation between TiP values from the same tissue but obtained from either GTEx or PostnatK. Pearson r and p-value are shown. **E.** Circle plot showing clustering of GTEx and PostnatK tissues based on computed TiP values. **F.** Violin plots of TiP value distributions for different tissues with samples from different gene expression resources as indicated and split into either protein-coding genes (PC) or neuron-specific genes (NS).

To quantify how preferential a given gene is expressed in a given tissue compared to all other tissues within the same dataset (i.e. GTEx, PrenatK, or PostnatK), we adopted the tissue-preferential (TiP) value computation, a z-score-based function, from Sonawane et al^13^. We modified this TiP value computation, allowing us to accommodate the highly variable and overall low number of samples per tissue in PrenatK and PostnatK. Similar to previous studies, we applied a TiP value cutoff of ≥ 2 within the GTEx dataset to call a gene significantly preferentially upregulated in a given tissue (Figure S1C)^7,13^. Due to much lower sample sizes in PrenatK and PostnatK compared to GTEx (Figure S1A), we lowered the TiP value cutoff for these datasets to match the same number of preferentially upregulated genes in GTEx when reduced to the four tissues brain, heart, kidney, and liver that are also present in PrenatK and PostnatK (GTEx-I, Figure 1C). The number of protein-coding (PC) genes detected as preferentially upregulated in at least one tissue was nearly 5,000 in GTEx and around 2,000 in both Kaessmann datasets (Table S1). In contrast, the number of genes not preferentially upregulated in any tissue (hereafter referred to as uniformly expressed genes) was approximately 9,000 in GTEx and about 12,000 in the Kaessmann datasets. Overall, we found good agreement between TiP values (Figure 1D, S1D) and sets of preferentially upregulated genes (Figure S1F) for the individual tissues when comparing GTEx with PostnatK, both representing gene expression levels of the postnatal stage. Brain, heart, and liver tissues from GTEx and PostnatK cluster together (Figure 1E), and all four PostnatK tissues show high Pearson correlation agreement with the corresponding GTEx tissue (Figure S1E). This indicates good comparability between GTEx and PostnatK and supports the joint use of all three datasets to study spatiotemporal patterns of preferential gene expression in pre- and postnatal stages across human tissues.

We further assessed to which extent the bulk RNA-seq data from GTEx, PostnatK, and PrenatK can resolve cell type-specific differential expression patterns. To this end, we assessed TiP values of cell type-specific genes for neurons, astrocytes, and oligodendrocytes in tissues from GTEx, PostnatK, and PrenatK in comparison to other PC genes (Table S2). We observed that neuron-specific genes exhibited significantly elevated TiP values in brain tissues across all three datasets, as well as in other neuronal and/or densely innervated tissues (Figure 1F). A similar, although less pronounced pattern was observed for astrocyte and oligodendrocyte genes (Figure S2). To further assess whether PrenatK reflects developmental stage-specific preferential expression of genes in the brain, we compared TiP values of neuronal stem cell (NSC) marker genes (Table S2) versus other PC genes in pre- and postnatal brain tissues. As expected, NSC marker genes showed TiP value distributions comparable to other PC genes in postnatal brain tissues. In the prenatal brain tissue, NSC marker genes displayed higher TiP values than other PC genes, consistent with their role in neurogenesis (Figure S1G). However, this difference did not reach statistical significance.

In conclusion, we established a robust RNA-seq processing protocol applicable to datasets of varying sizes and spatiotemporal resolutions. This enabled the generation of three comprehensive gene expression datasets (GTEx, PrenatK, and PostnatK) that jointly serve as a platform for investigating the spatiotemporal expression of any gene set of interest (Table S1). In the remainder of this study, we will leverage this resource to derive hypotheses regarding molecular mechanisms underlying brain-specific phenotypes.

### Tissue-preferential expression patterns of NDD-associated genes

We compiled a list of 983 genes that are associated with NDDs (hereafter referred to as NDD genes) based on whole exome sequencing studies from the Deciphering Developmental Disorders (DDD) consortium^9^ and the web resource Gene4Denovo^14^ (Figure S3A, Table S3). This enabled us to avoid particular bias towards highly studied genes. The vast majority of these genes are associated with undiagnosed developmental disorders, reflecting the fact that the majority of affected children remain without a molecular diagnosis due to limited understanding of the underlying molecular mechanisms (Figure S3B). Integrating this list of NDD-associated genes with GTEx TiP values, we find that less than 10% of NDD genes are preferentially upregulated in at least one of the three GTEx brain tissues. However, this fraction was significantly higher than that of other PC genes, as expected for a brain-related set of genes (Figure S3C). Repeating this analysis at the individual tissue level from GTEx, we find a significantly higher fraction of NDD genes to be preferentially upregulated in each of the three GTEx brain tissues as well as in heart (left ventricle) compared to other PC genes (Figure S3D). For the NDD genes that are significantly preferentially upregulated in the brain, we can postulate that they acquire brain-specific functions through brain-specific upregulation, likely explaining observed neurological manifestations when these genes are mutated. Nevertheless, the molecular mechanisms underlying brain-specific phenotypes for the remaining 90% of NDD genes that lack significant preferential brain upregulation remain elusive based on transcriptome data analysis.

We hypothesized that brain-specific functions of a subset of these genes may alternatively be driven by prenatal brain upregulation. Indeed, while the overall fraction of NDD genes preferentially upregulated in the brain remains stable between the pre- and postnatal stage, 15 NDD genes displayed prenatal brain preferential upregulation while not being significantly preferentially upregulated at the postnatal stage (Figure S3E). These genes include NEXMIF (neuronal migration and neurite extension factor), FOXP2 (transcriptional repressor which plays a role in neuronal differentiation), and SOX5, a well-known transcription factor expressed in neuronal stem cells (Figure S3E). Interestingly, we also identified NDD genes with known neuronal and synaptic functions (e.g., DLG4, KCNH1 and GABRA5) with the opposite preferential expression pattern, being preferentially upregulated in the brain postnatally but not prenatally (Figure S3E). However, while a subset of NDD genes shows preferential brain upregulation prenatally, postnatally, or consistently across pre- and postnatal stages of development, the majority of NDD genes are uniformly expressed across all three gene expression datasets (Figure S3F). Therefore, most NDD genes likely adopt brain-specific functions through mechanisms independent of upregulation in the brain at any developmental stage, thus highlighting the need for exploring alternative mechanisms.

To more systematically assess tissue-preferential expression patterns of NDD genes, we performed hierarchical divisive clustering (HDC) on the TiP values of all NDD genes across all GTEx tissues and identified 20 gene clusters (Figure 2A). Four of these clusters caught our attention based on their distinct preferential expression profiles. The first cluster largely overlapped with genes that we previously determined as significantly preferentially upregulated in the brain (hereafter referred to as BP cluster). Gene Ontology (GO) analysis revealed enrichment for ion channel function, synaptic transmission, synaptic membrane complexes as well as epilepsy and autism spectrum disorder (Figure 2B, Table S4). The second cluster of genes displayed preferential upregulation in muscle and heart tissues and showed enrichment for functions in mitochondrial energy production relating to the electron transport chain (ETC), the inner mitochondrial membrane (IMM) as well as cardiomyopathies and Leigh disease (Figure 2C, Table S4). We refer hereafter to this cluster as the IMM cluster. The third cluster of genes stands out for its preferential upregulation in kidney, liver, and adrenal gland and shows enrichments for mitochondrial matrix (MM) enzymes, organic acid catabolic processes as well as organic acidemia and amino acid metabolic disorder (hereafter referred to as MM cluster, Figure 2D, Table S4). The last cluster of genes to highlight consists of largely uniformly expressed genes (UE cluster) across all tissues. This cluster is enriched for genes encoding for proteins with DNA-binding functions such as transcription factors and histone modifying enzymes. It also shows enrichment for developmental disorder and intellectual disability (Figure 2E, Table S4). Finding clusters of NDD genes with preferential upregulation in tissues other than the brain (cluster 2 and 3) was rather surprising to us, triggering a more in depth analysis of these clusters as described below.

**Figure 2.**
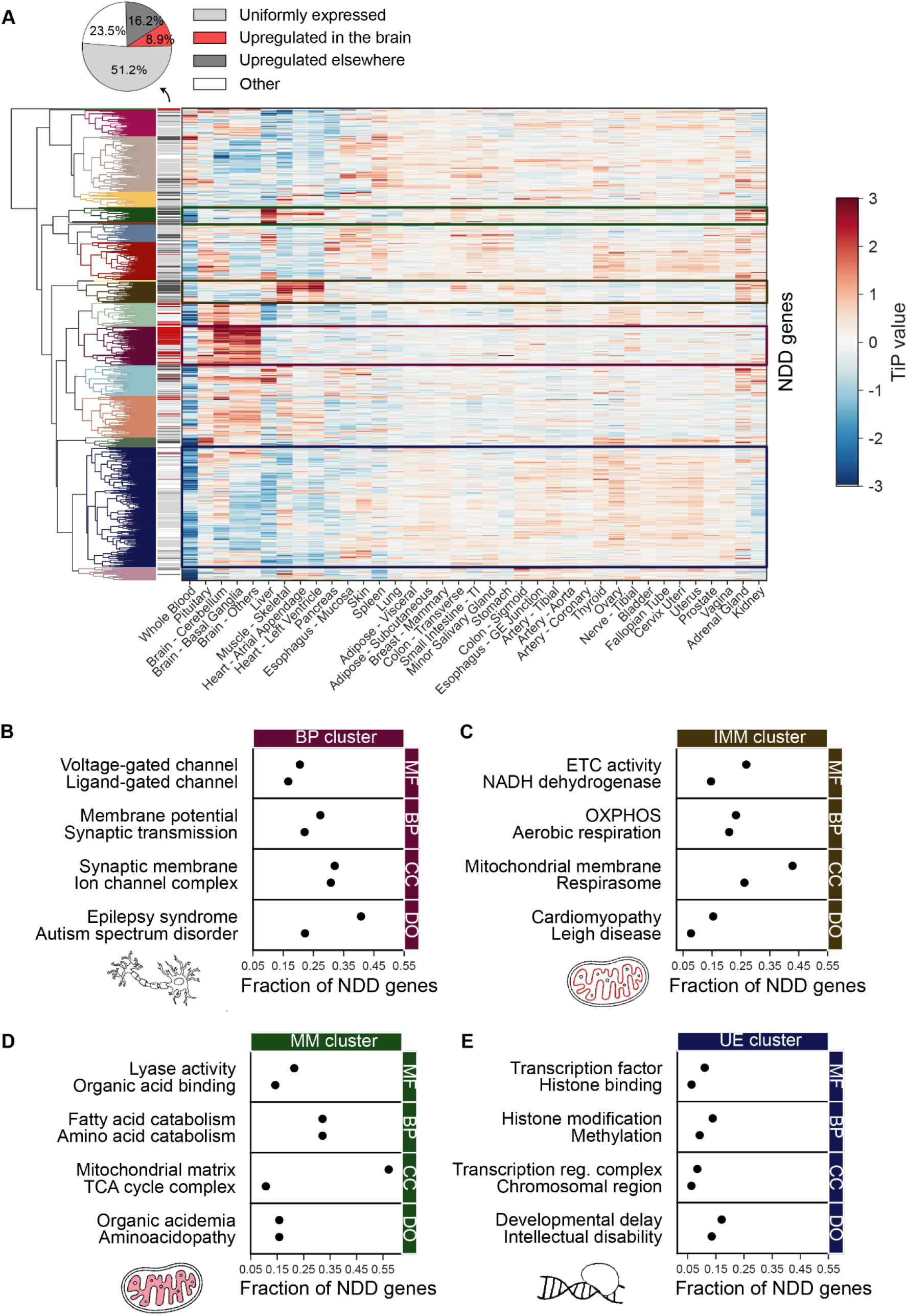
Exploration of clusters of NDD genes based on TiP values. **A.** Heatmap showing clustering of NDD genes based on TiP values across GTEx tissue samples and hierarchical divisive clustering. Four clusters selected for further inspection are indicated by frames. Classification of all NDD genes into uniformly expressed, upregulated in the brain or elsewhere, and others is based on a TiP value cutoff of 2. **B.** Enriched GO terms for NDD genes from the brain-preferential (BP) cluster as indicated in A. MF=molecular function, BP=biological process, CC=cellular component, DO=disease ontology. **C.** Enriched GO terms for NDD genes from the inner mitochondrial membrane (IMM) cluster as indicated in A. Abbreviations as in B. **D.** Enriched GO terms for NDD genes from the mitochondrial matrix (MM) cluster as indicated in A. Abbreviations as in B. **E.** Enriched GO terms for NDD genes from the uniformly expressed (UE) cluster as indicated in A. Abbreviations as in B.

### Mechanisms adapting mitochondria function to brain-specific contexts

To understand how preferential upregulation of NDD genes in tissues such as heart, skeletal muscle, kidney or adrenal gland is connected to neurodevelopmental defects caused by mutations in these genes we examined in more detail the NDD genes that are part of the IMM and MM cluster. We found that NDD genes within the IMM cluster correspond to components of nearly all complexes of the ETC, positioning them as direct contributors to mitochondrial ATP synthesis. This role aligns well with the previously mentioned enrichment of genes in this cluster for Leigh disease, a mitochondrial deficiency disorder arising from ETC failure^15^. In contrast, MM cluster genes predominantly encode Krebs cycle enzymes involved in β-fatty acid oxidation and amino acid catabolism. This pathway generates reducing equivalents (e.g., NADH and FADH₂) that feed electrons into the ETC and therefore, MM cluster genes represent a link between the cellular nutrient breakdown and mitochondrial energy production. Disruption of these genes drives organic acidemias, in which unmetabolized organic acids accumulate and lead to mitochondrial deficiency. Therefore, IMM and MM clusters converge on mitochondrial energy production and accordingly, their TiP profiles show preferential upregulation in high energy-demanding tissues, such as the heart, muscle, kidney and liver, but notably not in the brain. This is interesting, given that the brain is well established as one of the most energy consuming tissues in the human body^16^.

How else does brain tissue then meet its increased demand for ATP production if not through an upregulation of ETC and Krebs cycle proteins enabling an increase in mitochondria number? Neurons consume most of their energy at synapses and therefore depend on the transport of mitochondria along narrow axons to synaptic boutons, a process that requires coordinated mitochondrial fission and trafficking^17^. We hypothesized that, rather than upregulating ETC and Krebs cycle genes, neurons may instead upregulate genes that control mitochondrial transport, fusion, fission, cristae density regulation, and related processes, hereafter referred to as mitoregulatory genes. To test this, we curated a list of mitoregulatory genes from the literature and annotated them as either extra-mitoregulatory (EMR), if they localize outside of mitochondria, or intra-mitoregulatory (IMR), if they localize within the mitochondria but are involved in the regulation of mitochondrial function rather than energy production (Figure 3A). We further assembled a list of proteins that localize to the mitochondrial matrix (MM genes) and to the inner mitochondrial membrane (IMM genes) based on published experimental evidence^18^, both of which partially overlap with MM and IMM clusters of NDD genes (Table S5). We compared TiP value distributions for these four sets of genes across different energy-demanding tissues in GTEx and found that indeed MM, IMM, as well as IMR genes were more preferentially upregulated in heart, muscle, and adrenal gland compared to the remaining tissues (Figure 3B). In contrast, EMR genes exhibited significantly higher preferential upregulation in the brain compared to the remaining tissues (Figure 3B). This supports our hypothesis that the brain at least partially meets its specific energy demands via upregulation of genes that regulate mitochondrial function rather than by increasing mitochondria abundance and the expression of energy production-related genes. It also shows that tissue-preferential upregulation of genes is not necessarily a good indication for the relevance of these tissues for a given phenotype under study.

**Figure 3.**
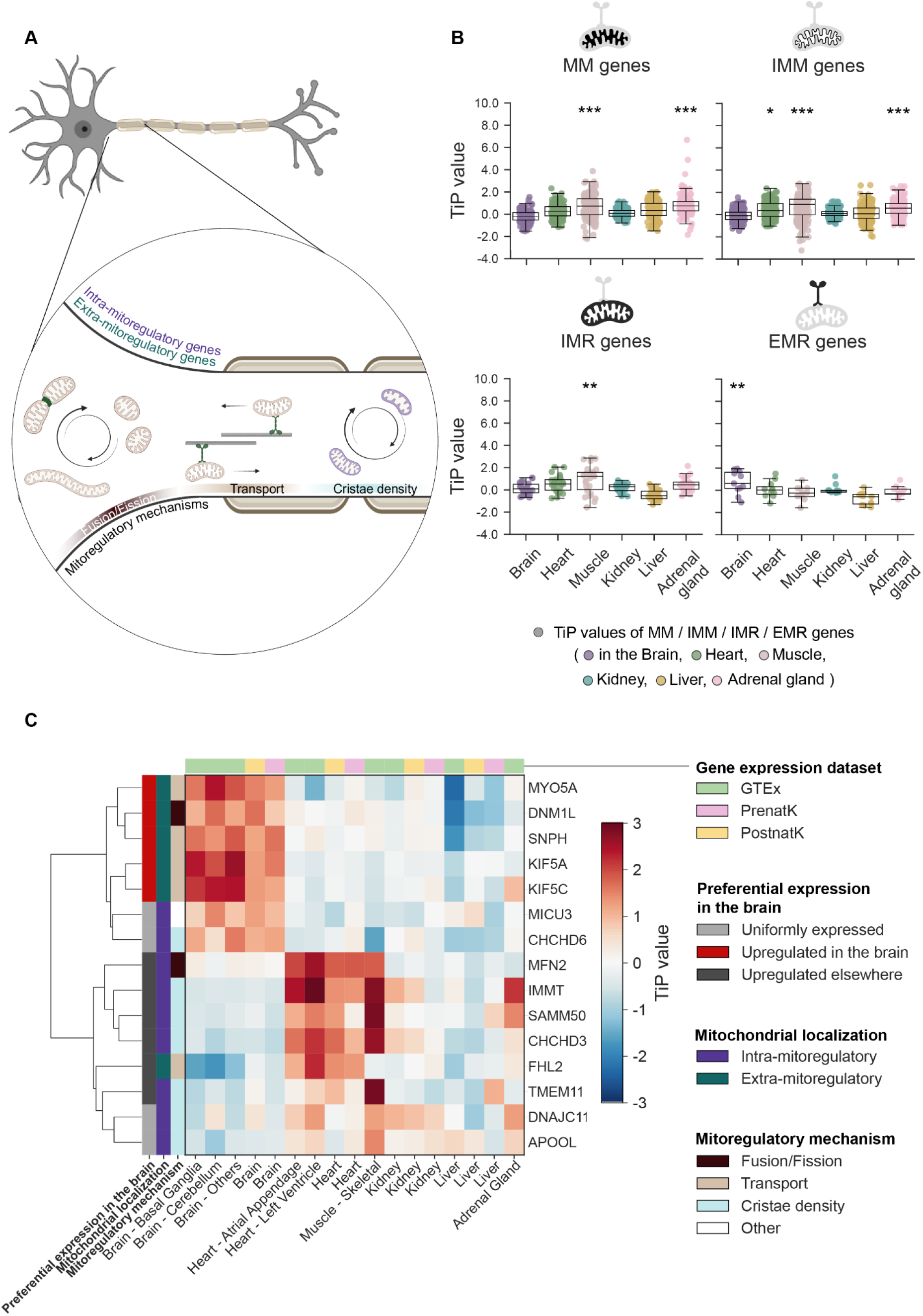
Exploration of preferential expression patterns across tissues of genes that regulate mitochondria function. **A.** Schematic of a neuronal cell and different processes that regulate mitochondria function. **B.** Box-whisker plots of TiP values from selected GTEx tissue samples and indicated groups of genes. MM=mitochondrial matrix, IMM=inner-mitochondrial membrane, IMR=inner-mitoregulatory, EMR=extra-mitoregulatory. **C.** Heatmap showing TiP values of selected genes indicated on the left functioning in mitochondria regulation across selected tissues from different expression datasets as indicated on the top and bottom and clustered based on hierarchical divisive clustering. Classification of genes into uniformly expressed, upregulated in the brain or elsewhere based on a TiP value cutoff of 2 in GTEx.

To further dissect preferential expression patterns of individual mitoregulatory genes, we analyzed TiP profiles for EMR and IMR genes across high energy-requiring tissues in GTEx, PrenatK, and PostnatK. This analysis revealed two coherent clusters (Figure 3C). The first cluster comprised genes that were significantly upregulated in the brain or showed the tendency towards brain-upregulation in GTEx, PrenatK or PostnatK. This cluster was dominated by EMR genes such as KIF4A, KIF5C, SNPH, MYO5A, and DNM1L. Interestingly, KIF4A, KIF5C, SNPH, and MYO5A mediate mitochondrial trafficking, while DNM1L controls mitochondrial fission, collectively indicating the importance of coordinated mitochondrial fragmentation and directed transport along axons^17^. Consistent with this, a few EMR genes are also associated with NDDs^19^, supporting the notion that disruption of mitochondrial positioning and dynamics may contribute to neurodevelopmental phenotypes. In contrast, the second cluster consisted of genes preferentially upregulated in other high energy-demanding tissues or genes that did not show significant preferential upregulation in any tissue (Figure 3C). This group largely corresponded to IMR genes, such as Mitofusin 2 (MFN2) - involved in mitochondrial fusion, and IMMT, member of the MICOS complex in the inner mitochondrial membrane, important for the maintenance of crista junctions. Their lack of brain-specific upregulation suggests that, even for mitoregulatory genes that localize within the mitochondria, increased expression is not the principal strategy neurons use to meet their metabolic requirements. Together, these findings suggest a model in which MM and IMM NDD genes confer brain-specific functions not through brain-preferential upregulation, but through their integration into brain-upregulated mitoregulatory pathways controlling mitochondrial dynamics, trafficking, and spatial organization.

### Brain-specific interaction partners of NDD genes

We next investigated potential mechanisms by which NDD genes that are uniformly expressed across tissues may nevertheless acquire brain-specific functions and contribute to neurodevelopmental phenotypes when mutated. To narrow the set of genes for mechanistic analysis, we re-clustered the UE subset of NDD genes using expression data from all GTEx, PrenatK, and PostnatK tissues and identified the largest subcluster, comprising approximately 100 genes, with consistent pre- and postnatal uniform expression (Table S6). This cluster was functionally and phenotypically representative of the broader UE NDD gene set, as evidenced by enrichment for similar GO terms, including DNA-binding functions, as well as shared enrichment for NDD-associated phenotypes such as intellectual disability (Table S7).

To next investigate whether the protein products of UE NDD genes adopt brain-specific functions via interaction with brain-upregulated proteins, we leveraged PPI data generated from high-throughput screens based on yeast two-hybrid (Y2H) and affinity purification coupled to mass spectrometry, hereafter referred to as the HuRI and BioPlex resource, respectively^7,20^. These resources were selected because they are not undersampling PPIs with tissue-specific proteins compared to other PPI resources such as those curated from the literature^7^. Furthermore, we wanted to focus on direct PPIs, i.e. proteins that are in direct physical contact with each other when binding, given that they are more amenable to mechanistic interpretation, particularly with respect to how mutations would affect protein binding. The Y2H-based resource HuRI predominantly contains direct PPIs^7^. In contrast, a substantial fraction of reported interactions in BioPlex are expected to be indirect^7^. To enrich BioPlex for likely direct PPIs, we restricted the dataset to interactions detected in both bait-prey orientations, yielding a subset of around 11,000 interactions, hereafter referred to as reciprocal BioPlex (BioPlex-R, Table S8). Using HuRI and BioPlex-R, we identified at least one PPI for 51 of the initial 100 UE NDD genes, resulting in a network of 512 interactions, most of which originated from HuRI. Of those, 22 UE NDD genes engaged in 45 PPIs with 36 interaction partners upregulated in the brain in GTEx, PrenatK or PostnatK, based on previously established cutoffs (Figure 4A, Table S9).

**Figure 4.**
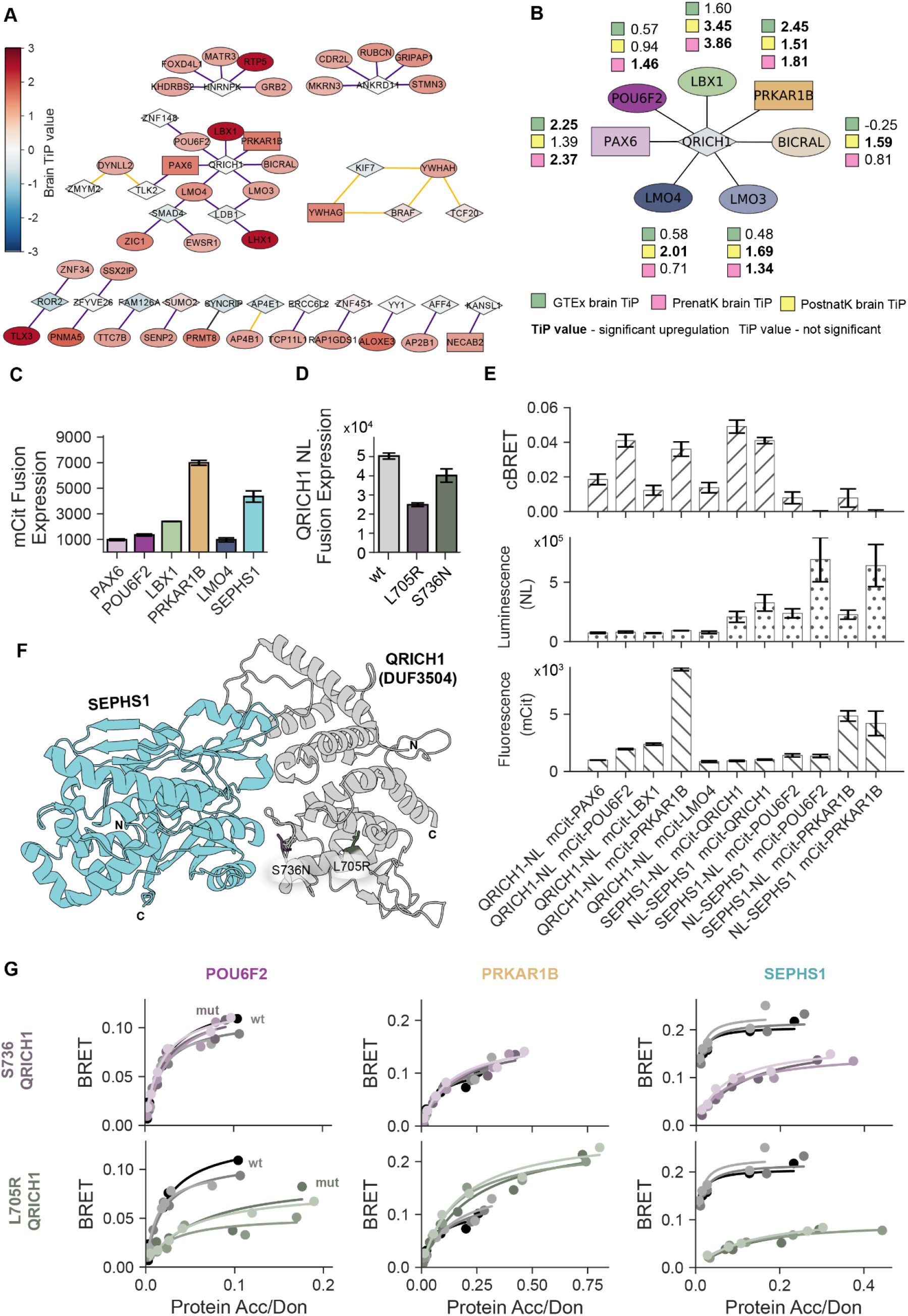
Brain-specific PPI analysis and experimental validation. **A.** Network showing proteins as nodes and physical interactions between them as edges. 22 uniformly expressed NDD genes with their brain-specific interaction partners are displayed. Nodes are colored according to the highest TiP value of the corresponding gene in any of the brain tissue datasets used in this study. Diamond nodes: uniformly expressed NDD genes; round nodes: non-NDD genes; square nodes: NDD gene and interactor of an NDD gene; orange edges: source BioPlex; purple edges: source HuRI; black edges: HuRI and BioPlex. **B.** Subnetwork from A. focussing on QRICH1 and interaction partners showing for each protein the TiP values from all three gene expression datasets for brain. **C.** Fluorescence values measured after transfecting cells with indicated proteins as mCitrine (mCit) fusions indicating their expression levels. Error bars indicate STD of three technical replicates. **D.** Total luminescence values measured after transfecting cells with QRICH1 wild type (wt) or indicated QRICH1 mutants as Nanoluc (NL) fusions indicating their expression levels. Error bars indicate STD of three technical replicates. **E.** Corrected BRET (cBRET) values as well as total luminescence and fluorescence values for indicated protein pairs. Error bars indicate STD of three technical replicates. **F.** Cartoon representation of a resolved structure showing the interaction between SEPHS1 and the DUF domain of QRICH1 (PDB code: 9HJT). The two tested QRICH1 mutants are labelled and drawn as sticks. **G.** BRET titration curves comparing QRICH1 wild type and mutants for their binding to three selected interaction partners as indicated on top of each graph. Each protein pair was measured in technical triplicates.

These 22 NDD genes are primarily associated with DNA-binding functions such as transcriptional regulation, as well as chromatin remodeling, histone modification, or DNA repair based on annotations in UniProt. Others are involved in mRNA processing. In contrast, the brain-upregulated interaction partners participate in neurogenesis, neuronal differentiation and fate specification, cell migration, and interneuronal junction formation. This supports a model in which uniformly expressed genes execute core cellular functions that are co-opted in the brain through interactions with brain-upregulated proteins, resulting in brain-specific functional modulation. One illustrative example of this model is an interaction cluster within our network involving the UE NDD genes BRAF, KIF7, and TCF20, and their brain-upregulated interaction partners YWHAH and YWHAG, the latter of which is itself implicated in neurodevelopmental disorders^21^. BRAF functions in MAPK signaling and cell proliferation regulation, KIF7 participates in primary cilium stability required for neuronal migration, axon elongation and neuronal connectivity, and TCF20 is a transcriptional co-regulator including the regulation of transcription factors functioning in neurodevelopment. On the other hand, YWHAH and YWHAG belong to the 14-3-3 (YWHA) family of proteins that modulate the subcellular localization and activity of proteins through phosphorylation-dependent binding, fine-tuning the function of their interaction partners in a cellular context-dependent manner^22^. Indeed, the interaction between YWHAG and BRAF has been previously well characterized and is known to regulate BRAF-driven cell proliferation^23^. Given the significant preferential upregulation of YWHAG in the brain identified in our study, this interaction may be particularly relevant for modulating BRAF signaling in the neuronal context.

Another interesting example is the interaction between the uniformly expressed NDD gene SYNCRIP and its brain-upregulated interaction partner PRMT8. SYNCRIP encodes an RNA-binding protein involved in mRNA processing and is known to bind mRNAs of genes involved in neuronal migration and outgrowth, regulating local translation in dendrites, and facilitating the axonal transport of mRNA of synaptic proteins such as synaptotagmin^24^. Thus, SYNCRIP’s neuron-specific functions likely at least partially arise from the binding of mRNA of brain-upregulated proteins. Furthermore, SYNCRIP’s RNA-binding activity is known to be regulated by arginine methylation, most notably through methylation by PRMT1^25^. In our interaction network, SYNCRIP was found to interact with PRMT8, another protein arginine methyltransferase that is preferentially upregulated in the brain and uniquely localizes to the cytoplasmic membrane. Given the dendritic localization of SYNCRIP, SYNCRIP and PRMT8 may co-localize in neurons, allowing PRMT8 to modulate SYNCRIP function in a brain-specific context through arginine methylation. Supporting this hypothesis, pathogenic NDD-associated variants have been identified both within the methyltransferase region of PRMT8 and within the region of SYNCRIP known to undergo arginine methylation, suggesting that disruption of this regulatory interaction may contribute to neurodevelopmental phenotypes.

The NDD gene with the highest number of brain-upregulated interaction partners in our interaction network was QRICH1 (Figure 4A, B). QRICH1 has been implicated in the regulation of gene expression, modulation of NF-κB signaling, proteostatic stress response, and the balance between cell survival and apoptosis, with outcomes that are highly dependent on cellular context^26–28^. More recently, QRICH1 has been reported to form the Zincore complex together with SEPHS1 regulating the expression of genes with roles in neurodevelopment^28^. However, given the uniform expression of QRICH1 and SEPHS1 across human tissues, the precise mechanisms leading to brain-specific gene expression regulation remain unclear. Pathogenic variants in QRICH1 are associated with the Ververi-Brady syndrome, a neurodevelopmental disorder with highly brain-specific manifestations, yet the mechanisms driving brain-specific phenotypes upon QRICH1 perturbation remain poorly understood^29^.

Six out of seven brain-upregulated interaction partners of QRICH1, which were all reported in HuRI, are involved in gene expression regulation, functioning as transcription factors, transcriptional co-regulators, or chromatin remodelers. PRKAR1B, the seventh partner protein, is a regulatory subunit of a cAMP-dependent protein kinase. Most of these proteins have known brain-specific functions, including the establishment of neuronal subtype identity and more generally, central nervous system development. To the best of our knowledge, interaction of QRICH1 with these partner proteins has not been reported outside of HuRI. QRICH1 has a particular protein architecture consisting of a small 50 residue long helical fold at the N-terminus, followed by an intrinsically disordered region of more than 400 residues that is highly glutamine-rich (hence its name), followed by a larger helical fold of 300 residues that is referred to as Domain of Unknown Function 3504 (DUF3504). Structural studies have revealed that the DUF mediates binding to SEPHS1^28^, which is reliably predicted by AlphaFold^30^ (note that the resolved structure of the SEPHS1-QRICH1 complex was released after AlphaFold’s training cutoff). However, exhaustive structural modelling attempts with AlphaFold-Multimer^30^ and AlphaFold3^31^ of QRICH1 with any of the brain-upregulated partner proteins did not result in any confident structural models precluding predictions of likely interacting regions between QRICH1 and its brain-specific partner proteins. We hypothesise that little co-evolutionary signal of likely more recently evolved tissue-specific PPIs might hinder successful structural modelling of these PPIs by AlphaFold.

To further experimentally support a likely functional relevance of these PPIs, we set out to first confirm binding of QRICH1 to these partner proteins using an orthogonal PPI assay that is based on bioluminescence resonance energy transfer (BRET)^32^. As part of this assay, two proteins of interest are exogenously expressed in a mammalian cell line as fusion constructs, one fused to a bioluminescent donor (nanoluc luciferase, NL) and the other to a fluorescent acceptor (mCitrine, mCit). The occurrence of a PPI between both proteins is determined by measuring energy transfer inside cells from an oxidized substrate of the donor to the acceptor. The efficiency of this energy transfer is highly distance dependent and can usually only occur if both proteins are directly interacting with each other. In addition to testing interactions between QRICH1 and its brain-upregulated partners, we also re-tested the interaction between QRICH1 and SEPHS1 as positive control. We obtained sequence-verified full length and expressed N-terminal mCit fusion constructs for PAX6, POU6F2, LBX1, PRKAR1B, LMO4, and SEPHS1, although expression of all partners apart from PRKAR1B and SEPHS1 was weak (Figure 4C). QRICH1 expression as NL or mCit fusion construct was also very weak (Figure 4D, E). We observed borderline interaction signals of QRICH1 with PAX6, LBX1 and LMO4, probably due to overall low expression levels. In contrast, interaction between QRICH1 and POU6F2 as well as with PRKAR1B yielded higher BRET signals as did the QRICH1-SEPHS1 interaction (Figure 4E, Table S10). Furthermore, we found no evidence for interaction between SEPHS1 and POU6F2 or PRKAR1B (Figure 4E). Of note, BRET signals are not directly comparable between PPIs because signal strength is directly dependent on the distance between donor and acceptor, which in turn is influenced by the size of the two interacting proteins, the binding affinity, and expression levels of both proteins.

To further evaluate if these PPIs are perturbed by NDD-associated mutations, we searched for QRICH1 variants that are unlikely to lead to a total loss-of-function. Thus, frameshift and truncating variants were excluded as well as missense variants that are buried within the DUF domain of QRICH1. We identified two interesting missense variants on the DUF domain: L705R, an uncharacterized variant, which is located a little further away from the interface with SEPHS1, and S736N, a likely pathogenic variant, which is close to SEPHS1 based on the resolved structure and found in the germline of two patients displaying Ververi-Brady syndrome^33^ (Figure 4F). Both residues are only partially exposed. L705R displayed a 2-fold reduction in QRICH1 protein levels compared to wild type while S736N remained almost as well expressed as determined from total luminescence signals measured in live cells after transient transfection with either wild type or mutant full length QRICH1 (Figure 4D).

Both variants clearly disrupted binding to SEPHS1 in BRET titration experiments, for which cells are transfected with increasing acceptor:donor plasmid ratios to monitor BRET signal saturation enabling BRET comparison at similar acceptor:donor protein ratios between wild type and mutant protein pairs. Interestingly, S736N had no detectable effect on QRICH1 binding by POU6F2 or PRKAR1B while L705R weakened binding to POU6F2 and potentially enhanced binding to PRKAR1B (Figure 4G). Unfortunately, BRET signals for the QRICH1 interactions with PAX6, LMO4, and LBX1 were too weak to clearly monitor interaction perturbation by QRICH1 variants.

Taken together, this data suggests that the L705R and S736N variants are likely deleterious to QRICH1 protein function via disrupted or altered PPIs with SEPHS1, POU6F2, PRKAR1B, and possibly additional QRICH1 interaction partners. These findings support a potential mechanism by which the uniformly expressed NDD gene QRICH1 acquires brain-specific functions by interacting with the brain-upregulated proteins POU6F2 and PRKAR1B, perturbation of which might cause neurodevelopmental phenotypes.

## Discussion

In this study, we established a computational pipeline to process and integrate spatiotemporal transcriptomic datasets, which enabled us to systematically analyze tissue-preferential gene expression across developmental stages. We applied this resource to predict molecular mechanisms that drive brain-specific phenotypes in neurodevelopmental disease contexts. Our results suggest that tissue-preferential gene expression rarely drives tissue-specific functions, as the majority of NDD-associated genes are not preferentially upregulated in the brain at neither prenatal nor postnatal stages. Instead, our results point to molecular context as a key determinant of tissue specificity as revealed by studying how the brain regulates mitochondria function and by exploring PPIs between uniformly expressed NDD genes and brain-specific partner proteins.

To quantify tissue-preferential expression, we computed TiP values, which capture the relative expression of a given gene in a given tissue compared to all other tissues^13^. This is highly advantageous over other approaches that use arbitrary expression level cutoffs to define tissue-specific expression^3,34^. We found that TiP value computation is less robust for datasets with small or highly unbalanced sample sizes. To overcome these limitations, we implemented downsampling and adapted the TiP value formula. However, defining comparable TiP value cutoffs across datasets remains a challenge. Due to low sample sizes and lack of discriminatory signal, we merged all pre- and postnatal developmental stages per tissue type in the Kaessmann dataset. We hope that with future efforts, gene expression datasets with increased sample sizes across tissues and developmental stages will become available enabling more fine-grained analysis of differential expression patterns of NDD genes. A further limitation in our study might have arisen from the use of bulk RNA-seq data, where gene expression signatures are averaged across different cell types within a given tissue, potentially masking genes that are preferentially expressed in rare or less abundant cell populations. Single cell RNA-seq data might be an interesting alternative in the future, should sufficiently broad and deeply sampled resources become available. Although RNA-seq data provide insight into transcript abundance across tissues, gene function is ultimately executed at the protein level, which were reported to only modestly correlate with mRNA levels^35^. Recently published spatiotemporal proteome resources promise a future extension of this work to systematically interrogate tissue-specific protein expression^36^.

Using our spatiotemporal gene expression resource, we confirm that most protein-coding genes are expressed across many tissues, often at comparable levels as reported previously^2,3^. Similarly, it has been reported that a substantial proportion of Mendelian disease genes is neither specifically nor preferentially expressed in the tissues primarily affected by their disruption^1,3^. Integrating spatiotemporal gene expression data with information on the molecular and functional context of genes, we revealed interesting alternative mechanisms for the acquisition of tissue-specific functions.

Regulatory roles of mitochondria in synapse function and neurogenesis are well established, and perturbation of mitochondria function, i.e. via genetic alterations, is associated with neurodevelopmental dysfunction^17,37^. Clustering of NDD genes by TiP values across human tissues revealed two clusters with upregulation of NDD genes in other tissues than the brain involving energy-demanding tissues such as skeletal muscle and heart. These NDD genes were enriched for functions relating to energy production and the ETC. By further exploring our gene expression resource we revealed patterns that suggest brain-specific upregulation of mitoregulatory but not of ETC genes in the brain to sustain brain-specific requirements of mitochondria transport, fusion, and fission. These processes ensure local energy production at the synapse^17,38^. Thus, our work provides complementary and data-driven support for the brain to meet its high energy demand not via upregulation of direct energy production as observed for heart and skeletal muscle but via upregulation of mitoregulatory processes. Similar analyses can be done systematically for other biological processes across tissues to further reveal mechanisms of tissue-specific adaptations.

The most direct way in which molecular context can alter the function of a gene is via direct interaction with partner proteins^3,6^. We systematically searched for possible brain-specific adaptations of uniformly expressed NDD genes via identification of interactions with brain-upregulated partner proteins. Using publicly available resources of PPI data, we uncovered a network of interactions between 22 uniformly expressed NDD genes and brain-upregulated partner proteins. For the NDD-associated protein QRICH1 we further provide experimental support for its interaction with the brain-upregulated partner proteins POU6F2 and PRKAR1B and disruption of these interactions by NDD-associated mutations. While these interactions require further experimental validation in neuronal cell or tissue models, to our knowledge, this work represents a first mechanistic hypothesis for how QRICH1 may acquire brain-specific functions and cause neurodevelopmental phenotypes upon perturbation, despite its uniform expression. Extension of this concept to explore cellular context-specific interaction with other biomolecules such as RNA and DNA promises to reveal additional putative tissue-adaptive mechanisms. Future efforts would equally benefit from available tissue or cell type-specific molecular interaction data, which, especially for PPIs, is still largely missing^39^.

Much of our work was focussed on the loss-of-function of NDD genes due to mutations and functional adaptation via interaction with tissue-specific partner proteins. However, tissue-specific phenotypes can also arise from tissue-specific gain of partner binding via gain-of-function mutations or via the tissue-specific absence of a paralog that therefore fails to compensate for the loss of an NDD-associated gene in that respective tissue^40^. We also neglected tissue-specific functional effects of genes that arise from alternative splicing, which is an especially wide-spread post-transcriptional regulatory mechanism in the brain^41^. The limiting factor here is short read RNA-seq data, which allows to quantify tissue-specific splice sites but not the description of entire tissue-specific transcripts. With advances in long-read RNA sequencing, these limitations might be surpassed in the near future.

We based our work on the assumption that most NDD-associated genes cause brain-specific phenotypes upon perturbation. It should be noted though that NDDs and other Mendelian disorders often do not display strictly tissue-specific phenotypes but rather affect multiple tissues in subclinical ways such as bone malformation or heart anomalies that frequently co-occur with developmental delay. More fine-grained phenotype annotations can be integrated in future work to improve the prediction of disease mechanisms. Despite these limitations we hope that our work inspires neuroscientists to explore new directions in the study of molecular mechanisms that drive neurodevelopment and others to extend this approach to different rare Mendelian-like disorders with tissue-specific manifestations.

## Materials and methods

### Processing of gene expression data

We downloaded the raw GTEx counts data and the raw Kaessmann bulk RNA-seq FASTQ data (GTEx release v8, accessed December 29, 2021; Kaessmann, accessed April 7, 2022)^2,11^. We performed read quality control and aligned the reads to the GRCh38 reference genome (release 26_GRCh38.p10) using STAR v2.7.3a^42^. Gene-level read counts were then quantified with RNA-SeQC v2.4.2^43^ using the GENCODE v26 collapsed gene model. The RNA-SeQC version used here includes a legacy mode reproducing the counting strategy of RNA-SeQC v1.1.9 used in GTEx v8. Next, cell line samples and testis were removed from GTEx, the latter because of its globally elevated gene expression profiles. In Kaessmann, testis was likewise excluded to match the exclusion criteria for GTEx, and ovary samples were additionally removed because they were only available at the prenatal and not postnatal stage. The transcriptomic similarity of samples was assessed using the “checkTissuesToMerge” function implemented in the YARN R package (v1.16.0)^12^, which applies multidimensional scaling (MDS) to identify tissues or subregions with similar expression profiles. Based on this analysis, the 13 GTEx brain subregions were collapsed into three groups: Basal Ganglia, Cerebellum, and Other. In the Kaessmann dataset, forebrain and hindbrain samples were merged into a single Brain category. MDS analysis also supported partitioning Kaessmann into prenatal (PrenatK) and postnatal (PostnatK) subsets, where PrenatK contained prenatal samples only and PostnatK contained neonatal and later postnatal samples. Following this, genes expressed at globally low levels were removed from GTEx, PrenatK, and PostnatK using the YARN function “filterLowGenes”. Genes were retained if they had at least one read in at least half of the samples within a given tissue group. Finally, read counts in each dataset were normalized using tissue-aware quantile normalization with the “normalizeTissueAware” function in YARN (method = qsmooth). PrenatK and PostnatK were normalized separately so that tissue-level differences were preserved within each dataset while allowing comparison of expression patterns between prenatal and postnatal stages.

### Computing tissue-preferential (TiP) scores

For each gene in each tissue, gene expression values were calculated as the median read count across all samples assigned to that tissue. Tissue-preferential expression (TiP) scores were then calculated using a modified version of the tissue-specificity metric described by Sonawane et al^13^. The original formulation was adapted to better accommodate datasets with fewer tissues and unequal sample representation across tissues, particularly the Kaessmann dataset, which after the processing, contains four tissues and contains disproportionately many samples for the brain compared to other tissues. The same modified metric was applied to both Kaessmann and GTEx for consistency:

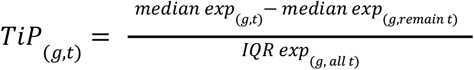

where *TiP* denotes the TiP value, *g* - gene, *t* - tissue, *median exp* - median expression (read count), *remain -* the remaining tissues, and *IQR exp -* the interquartile range, and *all* – all tissues.

To reduce the effect of brain overrepresentation in Kaessmann, brain samples were additionally downsampled by bootstrapping in PrenatK and PostnatK. This was done by generating 10 resampled datasets by randomly selecting n brain samples (without replacement), while retaining all non-brain samples. *n* was set to the mean sample number across non-brain tissues in the corresponding dataset: 12 for PrenatK and 35 for PostnatK. TiP scores were then calculated separately for each resampled dataset, and the median TiP value across the 10 iterations was used as the final TiP score for each gene-tissue pair in Kaessmann.

### Computing TiP score thresholds

Thresholds for gene expression and TiP values were defined by titration analysis. Genes with expression value ≥ 4 were considered expressed in a given tissue. To be considered preferentially upregulated in a given tissue in GTEx, a gene had to be expressed in that tissue, and additionally have a TiP score of ≥ 2 in that tissue. To be considered preferentially downregulated in a given GTEx tissue, a gene had to have a TiP score of ≤ -2 in that tissue, but still be expressed in the remaining tissues (median expression of ≥ 4 across the remaining tissues). Finally, to be labeled as uniformly expressed across all GTEx tissues in a given dataset, a gene had to be generally expressed across all tissues (median expression of ≥ 4 in each tissue in that dataset), and not have any significant TiP scores (all TiP scores ≥ -2 and ≤ 2).

For both Kaessmann datasets (PrenatK and PostnatK), the same gene expression threshold (≥ 4) was applied. To determine the significant TiP value thresholds for PrenatK and PostnatK, we implemented the following procedure. We restricted the GTEx dataset to the four tissues represented in the Kaessmann datasets (brain, heart, kidney, and liver), hereafter referred to as GTEx-I. For brain and heart, which include multiple subregions in GTEx, TiP values were averaged across the three brain, and two heart subregions for each gene, yielding a single TiP value per gene per tissue. The TiP value thresholds for PrenatK and PostnatK were then determined as the TiP values for which the fraction of preferentially expressed (up- and downregulated) genes matched the fraction of preferentially expressed genes in GTEx-I at the TiP value of +/- 2. This resulted in the TiP value cutoff of +/- 1.335 and +/- 1.405 for PrenatK and PostnatK, respectively.

### Generating a list of protein-coding (PC) genes

A list of PC genes was obtained from the Ensembl Human Genes database (release v104, accessed on 23/07/2021, downloaded from BioMart)^44^. The dataset was filtered to include only PC genes located on the primary assembly chromosomes (1-22), and genes from the mitochondrial chromosomes were excluded. The set of PC genes was further restricted to genes present in the preprocessed GTEx, PrenatK, and PostnatK datasets, yielding 15,664 unique ENSG IDs. PC genes with low expression across all tissues in GTEx, PrenatK, or PostnatK (expression values ≤ 4 in all tissues of a given dataset) were removed.

### Assessing GTEx and Kaessmann comparability

GTEx and Kaessmann comparability was first assessed by comparing the sets of preferentially upregulated genes in the four tissues present in GTEx and PostnatK. Given that the brain and the heart tissue is represented across multiple subregions in GTEx, a gene was considered upregulated in that tissue (the brain or the heart), if it was upregulated in any of the corresponding subregions. Next, we assessed the agreement between TiP values in GTEx-I and PostnatK by computing the Pearson correlation coefficient. For each gene-tissue pair, TiP values in GTEx-I were compared with the corresponding values in PostnatK. Correlation was calculated across all tissues collectively and subsequently for each of the four tissues individually. Next, pairwise similarity among GTEx and PostnatK tissues was quantified by computing the Pearson correlation coefficient for each GTEx-GTEx, GTEx-PostnatK and PostnatK-PostnatK tissue pair, using TiP values of all PC genes. The resulting correlation matrix was subjected to hierarchical clustering using the complete linkage method, as implemented in the clustermap function of the seaborn python package (v0.13.2)^45^.

### Obtaining lists of cell type-specific genes

Cell type-specific gene sets for neurons, astrocytes, and oligodendrocytes were derived from the meta-analysis by McKenzie et al^46^. Here, genes were evaluated using three complementary metrics of cell type-related expression: specificity, defined as expression restricted to one cell type relative to others; enrichment, defined as higher expression in one cell type compared with all remaining cell types; and absolute expression, defined as high expression in a given cell type irrespective of expression elsewhere. The data was obtained from Supplemental File 1 (“top_human_enrich”) of McKenzie et al^46^.

NSC marker genes were obtained from CellMarker (accessed October 18, 2023)^47^. CellMarker is a manually curated database of marker genes for more than 400 human cell types assembled from published studies and reference sources, including evidence from single-cell sequencing, flow cytometry, and immunostaining experiments. The NSC marker set provided in the CellMarker database and used in our analysis comprised 31 genes.

### Compiling a list of NDD genes

Publicly available data was obtained from the DDD study (DDD Consortium, *DDG2P_25_4_2021*; accessed April 26, 2021)^48^ and Gene4Denovo (v1.1, *Candidate_gene*; accessed April 18, 2021)^14^. From DDD, we retained genes annotated as *confirmed* in the *DDD category* field and that are associated with disorders showing a neurodevelopmental component, defined by organ specificity entries containing *Brain/Cognition*. From Gene4Denovo, we retained genes with FDR < 0.05 and relevant neurodevelopmental phenotype annotations in the *Groups* field, including *NDDs*, *ASD*, *ID*, *EE*, or *UDD*. Genes associated with noncoding variants were excluded (*noncoding* = 1). Deprecated gene symbols were updated to current HGNC-approved symbols. To this end, we retrieved the Ensembl Human Genes database (release v104, accessed on 23/07/2021, downloaded from BioMart)^44^, filtered for protein-coding genes on primary assembly chromosomes 1-22, and extracted Ensembl gene IDs, UniProt IDs, and HGNC symbols for mapping of the final NDD gene set.

### NDD gene clustering, GO and DO term enrichment analysis

NDD genes were clustered according to tissue-preferential expression profiles in GTEx (TiP values), using hierarchical divisive clustering (HDC) with complete linkage (method = “complete”), implemented in the seaborn (v0.13.2) python package (seaborn.clustermap)^45^. To determine the optimal cutting height of the resulting dendrogram, and define the number, size and composition of clusters of NDD genes, a range of cutting heights was evaluated. At each cut height, cluster membership was assigned with scipy.cluster.hierarchy.fcluster, and the resulting number of clusters and mean cluster size were evaluated across the dendrogram. Final cut heights were chosen to yield average cluster sizes suitable for downstream Gene Ontology (GO) and Disease Ontology (DO) enrichment analyses.

Cluster gene lists were stored as NumPy arrays and imported into R with reticulate (v1.42.0)^49^. For each cluster, GO enrichment was tested across the biological process (BP), cellular component (CC), and molecular function (MF) ontologies using enrichGO from clusterProfiler (v4.4.4)^50^, and DO enrichment was tested using enrichDO from DOSE (v3.22.1)^51^. Human gene annotations were provided by org.Hs.eg.db (v3.21.0)^52^. For each cluster and enrichment analysis, input genes were tested against a background of all genes with available GO or DO annotation. Multiple testing correction was performed using the Benjamini-Hochberg method, and adjusted *p* values < 0.05 were considered significant. Only gene sets containing at least five genes were analyzed.

### Compiling lists of mitochondrial and mitoregulatory genes

Genes that regulate the functionality of mitochondria (mitoregulatory genes) were compiled from López-Doménech and Ralf Kittler^17^. Gene lists were extracted and annotated according to functional category and subcellular localization based on this study. Functional annotations were simplified into five categories: mitochondrial transport, mitochondrial fusion and fission, cristae organization, mitochondrial degradation, and energy production. In addition, genes were classified by localization as extramitochondrial or intramitochondrial. Extramitoregulatory (EMR) genes encode proteins that do not reside within the mitochondria but associate with or regulate the organelle from outside, whereas intramitoregulatory (IMR) genes encode proteins localized to mitochondrial compartments (including the outer membrane, inner membrane, intermembrane space, or matrix).

As external mitochondrial reference sets, we also compiled genes annotated to the mitochondrial matrix (MM) and inner mitochondrial membrane (IMM). MM and IMM gene lists were retrieved from UniProt using the advanced search interface with the following filters: taxonomy restricted to *Homo sapiens* (9606), subcellular location term set to *Mitochondrion matrix* [SL-0170] for MM genes or *Mitochondrion inner membrane* [SL-0168] for IMM genes, evidence restricted to experimental, and reviewed entries only (Swiss-Prot). This yielded 150 MM genes and 189 IMM genes (accessed May 5, 2025; downloaded from UniProt)^18^.

### UE NDD gene clustering, GO and DO term enrichment analysis

Clustering and GO/DO term enrichment analyses were performed as described earlier. From clustering all NDD genes (see above), we obtained one cluster comprising 242 genes, which exhibited uniform expression across all GTEx tissues (UE genes). Here, this cluster was subjected to a second round of clustering to identify subclusters. For this purpose, gene expression data from GTEx, as well as PrenatK and PostnatK tissues, were included. For the identified sub-clusters we computed uniform expression scores, based on the relative abundance of non-TiP genes across GTEx, PrenatK, and PostnatK in each cluster. For each cluster in each of the three gene expression datasets, genes were classified as brain-TiP (significantly preferentially upregulated in the brain) or non-TiP (not significantly preferentially upregulated in the brain), and the ratio of TiP to non-TiP genes was calculated. These three dataset-specific ratios were then averaged. The final score was computed as the natural logarithm of cluster size minus the natural logarithm of the mean ratio, with a constant of 0.01 added to the denominator term to avoid undefined values:

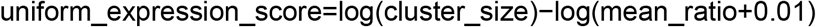

Higher scores correspond to larger clusters with a lower proportion of brain-TiP genes. Based on this, a cluster of 100 UE NDD genes was identified among the eight clusters, and was used in downstream analyses.

### Retrieval and processing of protein interaction data

HuRI data was downloaded from the HI-union release (accessed September 1, 2021; https://interactome-atlas.org/download, “HI-union”)^53^ and further filtered to remove homodimeric interactions, yielding 63,242 PPIs represented by ENSG identifiers. BioPlex data were obtained from BioPlex 3.0 interactions in 293T cells (release v3.0, accessed July 27, 2021, https://bioplex.hms.harvard.edu/interactions.php, “BioPlex 3.0 Interactions (293T Cells)”)^20^. This dataset was additionally processed by removing homodimers. The resulting processed BioPlex dataset contained 114,269 PPIs represented by ENSG IDs. Using data provided to us by the BioPlex authors directly, we further filtered BioPlex to gene pairs detected in both bait-prey orientations, yielding 11,456 reciprocal interactions (BioPlex-Reciprocal, “BioPlex-R”).

### Cell line culture and maintenance

HEK293 cells were originally obtained from DSMZ (catalog number ACC305). Cells were cultured in DMEM (Thermo Fisher Scientific, 21969035), supplemented with 10% FBS (Thermo Fisher Scientific, A5256701),1% L-Glutamin (Thermo Fisher Scientific, 25030024) and 1% Penicillin/Streptomycin (Thermo Fisher Scientific, 15140122). Cultures were kept at 37°C with 85% humidity and 5% CO₂. Routine passaging was performed every 2-3 days using 1mL of 0.05% trypsin (gibco, 25300-054, Thermo Fisher Scientific, 25300054), with 1,0 × 10⁶ cells seeded per T25 flask (Sarstedt, 83.3910.002). The purchased cell line was initially expanded, allotted and frozen at -150°C for storage. For quality control, cultures were screened for mycoplasma contamination before freezing and at certain intervals using the mycoplasmacheck from Eurofins. No additional authentication of the cell line was performed.

### Plasmid construction

The donor vector pcDNA3.1-myc-NL-GW (Addgene ID #113446), pcDNA3.1 GW-NL-cmyc (Addgene ID #113447) and control plasmids pcDNA3.1(+), pcDNA3.1 NL (Addgene ID #113442), and pcDNA3.1-PA-mCit (Addgene ID #113443) were kindly provided by the Wanker group (Max Delbrück Center for Molecular Medicine, Germany). The acceptor vector pcDNA3.1-mCit-His3C-GW and pcDNA3.1 GW-His3C-mCit were generated in house.

### GATEWAY cloning procedure

Cloning was done as described previously with some modifications^54^. Full-length wild type human open reading frames (ORFs) cloned in GATEWAY-compatible entry vectors (ORFeome Collaboration, 2016)^55^ were maintained as glycerol stock. A sterile swab from each bacterial glycerol stock containing human ORFs was used to inoculate individual wells of 96-well plates (Thermo Fisher Scientific, 163320), each containing 200 µl of LB medium supplemented with appropriate antibiotics. To minimize evaporation during incubation, the plates were sealed with sterile self-adhesive airpore film (Fisher Scientific, 10642674). Cultures were grown overnight at 37 °C with shaking at 190 rpm. The inoculated ORF cultures were used directly as template for PCR amplification. In PCR stripes (Starlab, I1402-3700) 4µl of the inoculated culture was used per each 50µl PCR reaction using Phusion High-Fidelity DNA Polymerase (NEB, M0530S), and primers annealing to vector flanking regions (forward: 5′-TTGTAAAACGACGGCCAGTC; reverse: 5′-GCCAGGAAACAGCTATGACC). Thermal cycling was carried out at 98 °C for 10 s, 55 °C for 30 s, and 72 °C for 3 min for 30 cycles. PCR products (5 µl per well) were analyzed on 1% agarose gels containing Ethidium Bromide. Each well was loaded with 1 µl of 6x gel loading dye purple (NEB,B7024A), and product sizes were assessed using 5 µl of 1kbPlus DNA ladder (NEB,N3200S). For downstream cloning, 1 µl of each amplified PCR product was mixed with 200 ng of the above-mentioned destination vectors in a 10 µl Gateway LR recombination reaction using 4× LR Clonase enzyme mix (Thermo Fisher Scientific, 11791020), performed in a 96-well PCR plate and incubated overnight at 25°C to generate the final expression constructs. For increased efficiency 1µl Proteinase K (2µg/µl) was added after the LR reaction and incubated for 10 min at 37°C. The entire 10 µl LR reaction mixture was subsequently transformed into 30 µl of chemically competent E. coli DH5α cells (Thermo Fisher Scientific, 18265017) in a 96-well PCR plate. Following transformation, cells were recovered in 80 µl of SOC medium at 37 °C for 1 h without shaking. A total of 70 µl from each transformation was plated onto square 48-well LB agar plates and incubated overnight at 37 °C to allow colony formation. Individual colonies were manually picked and cultured in 96-deep-well plates (Starlab, E2896-2110) containing 2 ml of LB medium supplemented with 100 µg/ml ampicillin (Sigma Aldrich, A9518). Cultures were incubated at 37 °C with shaking at 700 rpm for 24 h in an IncuMixer. Plasmid DNA was extracted using the QIAprep Spin Miniprep kit 250 (Qiagen, 27106) following the manufacturer’s protocol. DNA concentrations were determined using a Nanophotometer, and samples were diluted to a final concentration of 100 ng/µl. For sequence validation, 600 ng of plasmid DNA was subjected to full-length Sanger sequencing using backbone-specific primers (NanoLuc forward: 5′-GAACGGCAACAAAATTATCGAC or mCitrine forward: 5′-AGCAGAATACGCCCATCG with pEXP reverse: 5′-GGCAACTAGAAGGCACAGTC and NanoLuc reverse: 5′-CCCCGAGATTCTGAAACAAAC or mCitrine reverse: 5′-AGTGTGAGTTTCCCATATGTCG with pEXP forward: 5′-GGCTAACTAGAGAACCCACTG) and ORF-specific primers (Table S12). All sequence-validated ORFs utilized in this study along with their sequences are listed in Table S11.

### Site-directed mutagenesis

Mutagenesis was done as described previously with some modifications^54^. The site-directed mutagenesis was done with the Q5 Site-Directed Mutagenesis Kit (NEB, E0554S). Primers were designed manually following the guidelines from NEB. Site-directed mutagenesis was performed in PCR stripes using 10ng of plasmid DNA template and primer pairs in 50µl reactions. Amplification was carried out using Phusion High-Fidelity DNA Polymerase (NEB, M0530S) under the following conditions: initial denaturation at 98 °C for 2 min; 25 cycles of 98 °C for 15 s, annealing (temperature optimized per primer pair) for 15 s, and extension at 72 °C for 5 min. To remove template DNA, 1 µl of DpnI (NEB, R0176S) was added directly to each reaction and incubated at 37 °C for 1 h, followed by enzyme inactivation at 65 °C for 20 min. PCR products (6 µl) were confirmed on agarose gels and size verification was performed using 6µl 1kb Plus DNA Ladder (NEB, N3200S). After confirming the PCR products, KLD reaction from the Q5 site directed mutagenesis kit was performed using 1µl PCR product, KLD buffer and KLD enzyme mix in a 10µl reaction volume, mixed up and down and incubated for 5min at room temperature. Following Chemically competent E. coli DH5α cells (30 µl) were transformed with 3 µl of the KLD reaction mix. Cells were recovered in 800µl SOC medium at 37°C for 1h at 350 rpm. Subsequently recovery, 70µl of each transformation was plated onto ampicillin agar plates and incubated overnight at 37°C. Individual colonies were manually selected and grown in 5ml round bottom tubes containing 5ml LB, supplemented with 100µg/ml ampicillin. Cultures were incubated overnight at 37°C with shaking at 190rpm. Plasmid DNA was isolated using the QIAprep Spin Miniprep kit 250 (Qiagen, 27106), quantified using a nanophotometer and sent for Sanger sequencing.

### Sequencing

Sanger sequencing was performed in PCR tubes (Starlab, I1402-3700) to validate the full insert. For each sample, 400–600 ng of plasmid DNA was used together with tag- or ORF-specific primers (Table S12) to ensure complete coverage of the open reading frame and engineered mutation sites. For each reaction, 1 µl of the appropriate sequencing primer was added. Subsequently, 6 µl of diluted plasmid DNA was added and sent for Sanger sequencing to StarSEQ. Upon return of sequencing data, reads were manually checked with Benchling. Sequences of wild type and mutated inserts are available in Table S11.

### BRET assay

The BRET assay was conducted as described previously with the following modifications^54^.

### Transfection

HEK293 cells were cultured in high-glucose (4.5 g/l) DMEM (Thermo Fisher Scientific, 21969035), supplemented with 10% FBS (Thermo Fisher Scientific, A5256701), 1% L-Glutamin (Thermo Fisher Scientific, 25030024) and 1% Penicillin/Streptomycin (Thermo Fisher Scientific, 15140122). Cells were maintained at 37 °C with 5% CO₂ and 85% relative humidity and subcultured every 2–3 days. For BRET assays, reverse transfection was performed using Lipofectamine 2000 (Thermo Fisher Scientific, 11668019) in Opti-MEM (Thermo Fisher Scientific, 51985034) according to the manufacturer’s protocol. Cells were seeded in 96-well white flat-bottom plates (VWR, 392-0229) at a density of 4.0 × 10⁴ cells per well in phenol red-free, high-glucose DMEM (Thermo Fisher Scientific, 21063029) supplemented with 5% FBS (Thermo Fisher Scientific, A5256701) and 1% Pen/Strep. Transfections were carried out with a total of 200 ng plasmid DNA per well. When the DNA amount of expression plasmids was below 200 ng, pcDNA3.1(+) was added as carrier DNA to reach the total amount. Prior to transfection, plate layouts were designed including appropriate controls: NL-stop + pcDNA, mCit-stop + pcDNA, pcDNA-only wells, and wells with untransfected cells. Control DNAs (PA-mCit-Stop, NL-Stop, pcDNA3.1) were prepared in PCR strips (Starlab, I1402-3700) for efficient pipetting. NL constructs were further diluted to 4 ng/µl. Cells were harvested, trypsinized, counted, centrifuged, and adjusted to a final concentration of 2.67 × 10⁵ cells/ml in phenol red-free DMEM and 150µl of this cell suspension was pipetted into each well. Plates with the transfected cells were incubated at 37 °C with 5% CO₂ and 85% relative humidity for 48 hours prior to measurement.

### Single point BRET measurements

All measurements were performed using an Infinite M200 Pro microplate reader (Tecan). First, 100 µl of the medium were aspirated from each well. mCitrine fluorescence (FL) was measured in intact cells using excitation/emission wavelengths of 513/548 nm, a gain setting of 100, and 4 point measurements per well, which were averaged to represent the FL value for this well. Following fluorescence measurement, coelenterazine-h (PJK Biotech GmbH, 102181) was added to each well at a final concentration of 5 µM. The plate was briefly shaken for 15 s and incubated for 15 min inside the plate reader at 37 °C. After incubation, total luminescence (totLu) was measured per well, followed by short-wavelength and long-wavelength luminescence using the BLUE1 (370–480 nm) and GREEN1 (520–570 nm) dual filter setting, respectively, with an integration time of 1000 ms. Transfections with NL-stop at 2 ng in one well were used to determine the bleedthrough. Transfections with pcDNA3.1 at 200 ng in one well were used to determine background totLu and FL signals. Single point protein interaction measurements were done with DNA ratios for transfection of 20:50ng for pairs with QRICH1-NL fusions and with 2:50ng for pairs with SEPHS1-NL and NL-SEPHS1 fusions. Corrected BRETs (cBRETs) for these single point measurements were obtained by subtracting from the BRET of the protein pair the max BRET of two control measurements obtained from NL-STOP with proteinB mCit fusion and proteinA NL fusion with mCit-Stop.

### BRET titrations

For each protein pair the following six different donor:acceptor DNA ratios were used for transfections: 20:12.5 ng, 20:25 ng, 20:50 ng, 20:100 ng, 10:100 ng, and 10:200 ng. This is different from previous procedures^54^ because of very low QRICH1 expression levels as NL fusion. Each well contained 200 ng of DNA in total, with pcDNA3.1(+) added to the well to reach this amount. DNA mixtures for each protein pair were first prepared as master stocks in a 96-well PCR plate for three technical replicates. From this master plate, the DNA mixtures were distributed into three 96-well white flat-bottom plates (VWR, 392-0229). Transfection and subsequent measurements were conducted as described above.

Average background FL and totLu signals (see above) from one biological replicate were subtracted from each measured FL and totLu value, respectively. The average bleedthrough (see above) from an experiment was subtracted from the BRET signals. After background and bleedthrough correction all negative FL, totLu, and BRET values were set to zero. For every protein pair six titration points per technical replicate were measured (see above). Titration points were excluded, if totLu < 15000. Titration points were fitted, if at least five titration points for a given technical replicate passed this cutoff and at least three of these titration points had FL values ≥ 300. These filtering criteria were put in place to ensure that only titration points from measurements with sufficient donor (totLu) and acceptor (FL) expression levels were used to derive a fit. The leastsq function from the scipy.optimize python package was used to fit the remaining titration points per replicate to a simple 1:1 binding model with BRET = ((A/D) * BRETmax)/(BRET50 + (A/D)) where A = acceptor protein levels (FL) and D = donor protein levels (totLu).

## Acknowledgements

We thank all members of the Luck lab for helpful discussions throughout the study. We thank the protein production, media and IT core facilities from IMB. We thank Edward Huttlin for providing us with the BioPlex-R dataset. The research performed in this study was funded by the Deutsche Forschungsgemeinschaft (DFG, German Research Foundation) – Project-ID 449991970 awarded to K.L. This study makes use of data generated by the DECIPHER community. A full list of centres who contributed to the generation of the data is available from https://deciphergenomics.org/about/stats and via email from. DECIPHER is hosted by EMBL-EBI and funding for the DECIPHER project was provided by the Wellcome Trust (grant number WT223718/Z/21/Z).

## Use of AI

AI has been used to improve the style of selected paragraphs in the abstract and introduction.

## Code and data availability

All code relevant to this work is available on github (github.com/KatjaLuckLab/NDD_gene_expr_manuscript). Data is available as supplementary material as part of this manuscript.

## Author contributions

K.L. designed the study, M.D., S.K. and F.K. performed computational work, M.W. performed experiments, K.L. and M.D. performed experimental data analysis, M.D., K.L., M.W. and S.K. wrote the manuscript, K.L. provided the funding.

## Supplementary figures

**Figure S1.**
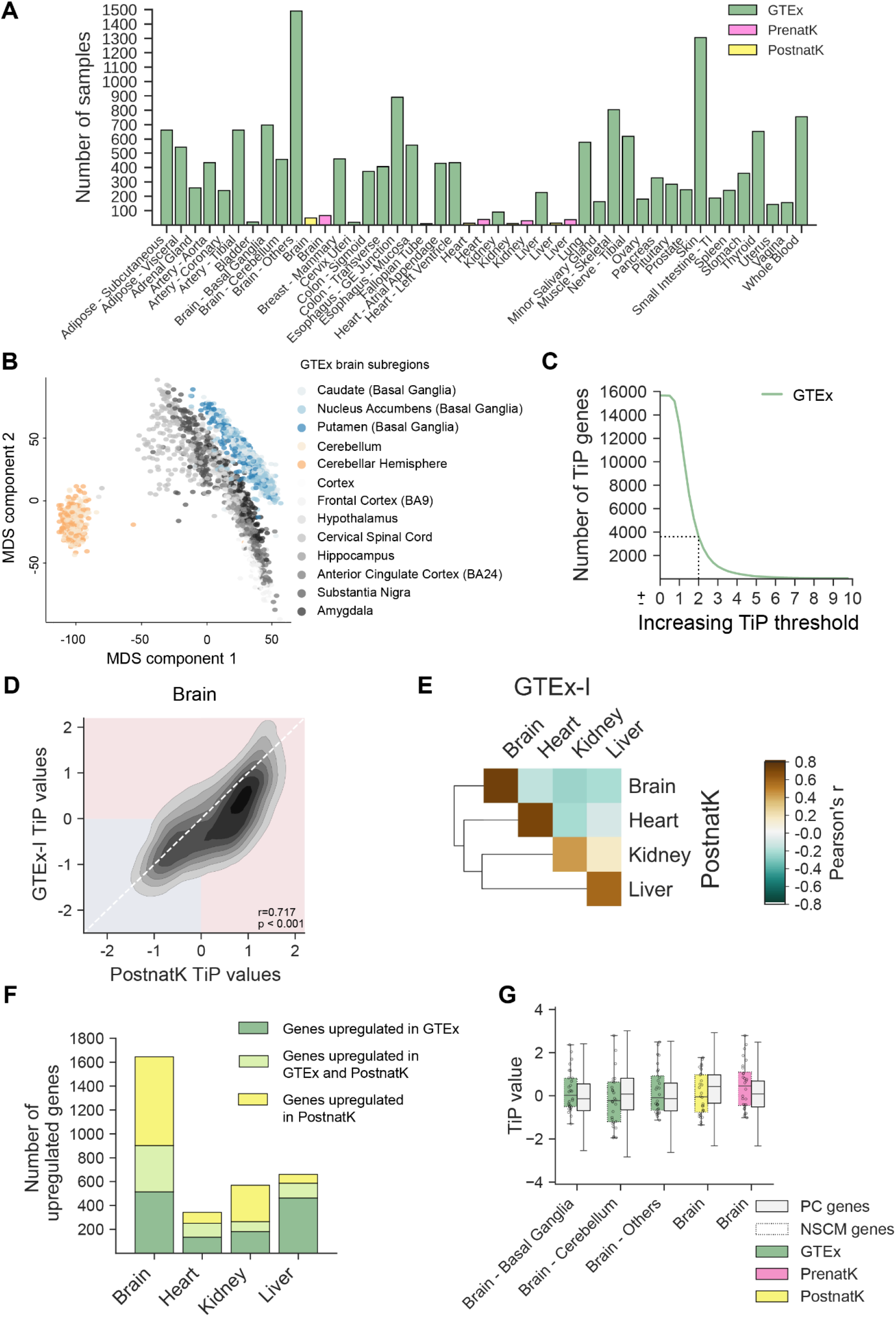
Characterization and comparison of gene expression datasets. **A.** Bar graph showing the number of samples for each of the tissues and developmental stages from all three gene expression datasets used in this study, as indicated. **B.** MDS plot showing how all 16 brain-subregions from GTEx cluster based on RNA expression values cluster into three distinct groups. **C.** Titration plot showing the decreasing number of TiP genes in GTEx for increasing TiP value cutoffs. Dashed lines indicate the chosen cutoff of 2. **D.** Density plot of TiP values of genes from brain tissue from GTEx-I and PostnatK. Pearson r and p-value are indicated in the plot. **E.** Heatmap showing the Pearson r for TiP values from pairs of tissues from GTEx-I and PostnatK. **F.** Bar graph showing the number of TiP genes per tissue for GTEx and PostnatK indicating the number of genes that are only TiP in one or both datasets. **G.** Box whisker plot showing the distribution of TiP values of neuronal stem cell marker (NSCM) genes versus protein-coding (PC) genes for different brain samples.

**Figure S2.**
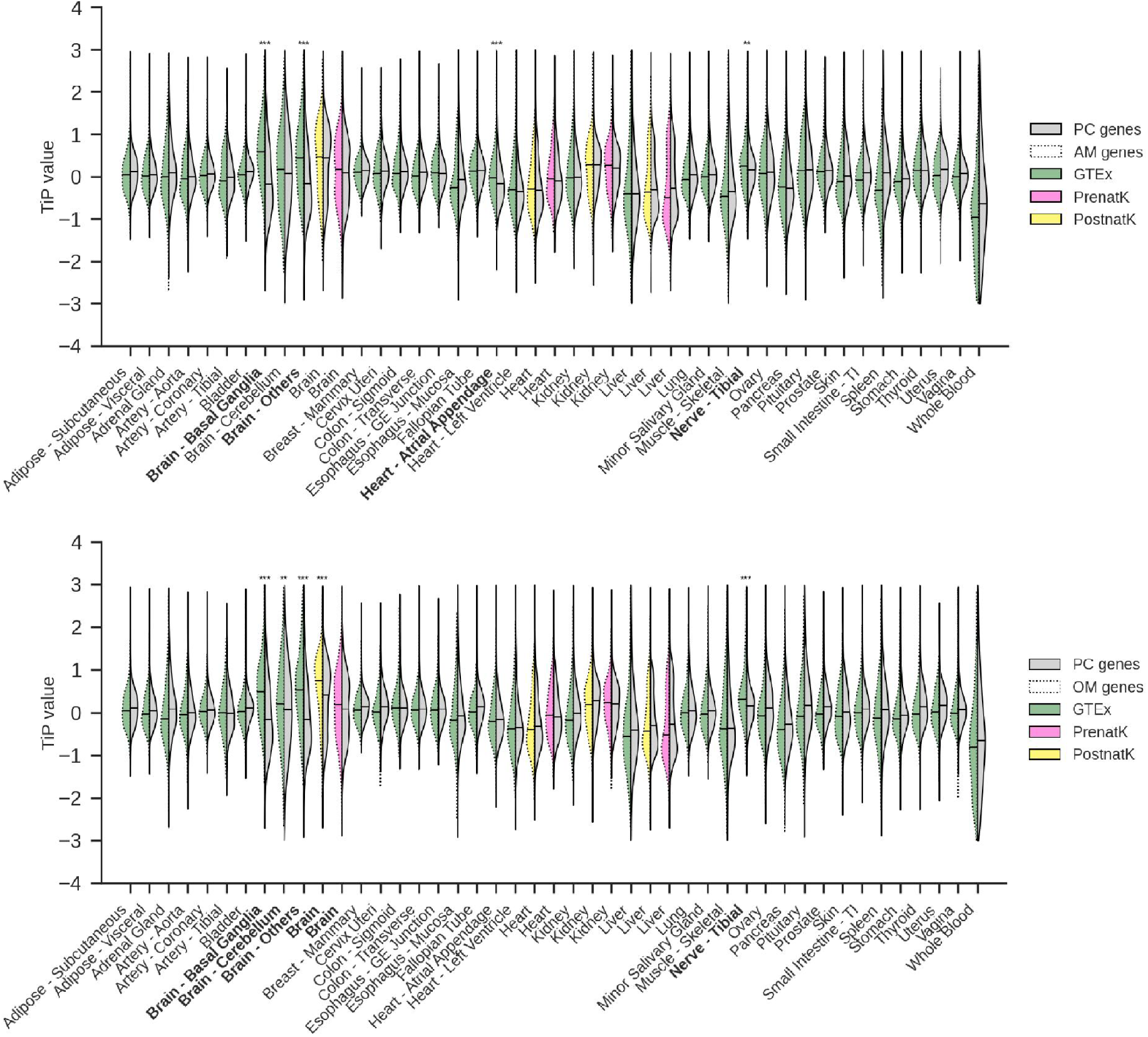
Violin plots of TiP value distributions for different tissues with samples from different gene expression resources as indicated and split into either protein-coding genes (PC) and astrocyte marker genes (AM, top) or oligodendrocyte marker genes (OM, bottom).

**Figure S3.**
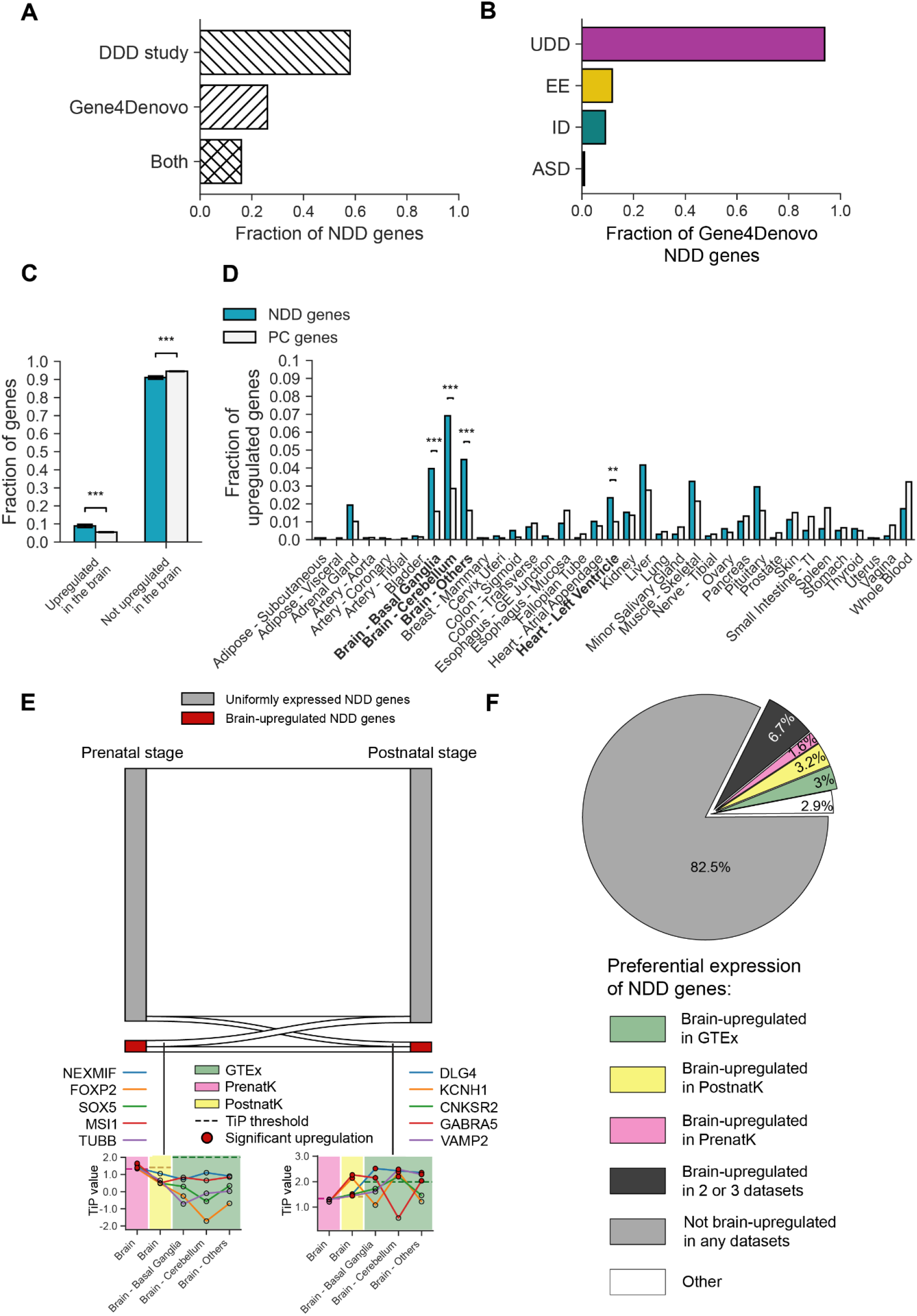
Characterization and preferential expression analysis of NDD genes. **A.** Bar plot showing the fraction of NDD genes per source. **B.** Bar plot showing the fraction of NDD genes from Gene4Denovo per disease subtype. UDD=undiagnosed developmental disorder, EE=epileptic encephalopathy, ID=intellectual disability, ASD=autism spectrum disorder. **C.** Bar plot showing the fraction of genes split into NDD genes and protein-coding (PC) genes and if they are significantly upregulated in the brain based on GTEx data and a TiP value cutoff of 2 or not. **D.** Bar plot showing the fraction of genes of either NDD genes or PC genes per tissue from GTEx that are significantly upregulated. Tissues for which both gene sets are significantly different are indicated with asterisks. **E.** Top: sankey plot showing the flow of NDD genes between brain pre- and postnatal stage from Kaessmann split into if they are considered upregulated in the brain or uniformly expressed. Bottom: dot plots showing TiP values for different brain samples as indicated and selected marker genes that change preferential expression status between the pre- and postnatal developmental stage. **F.** Pie chart showing fractions of NDD genes based on their upregulation in brain-related tissues.

**Figure S4.**
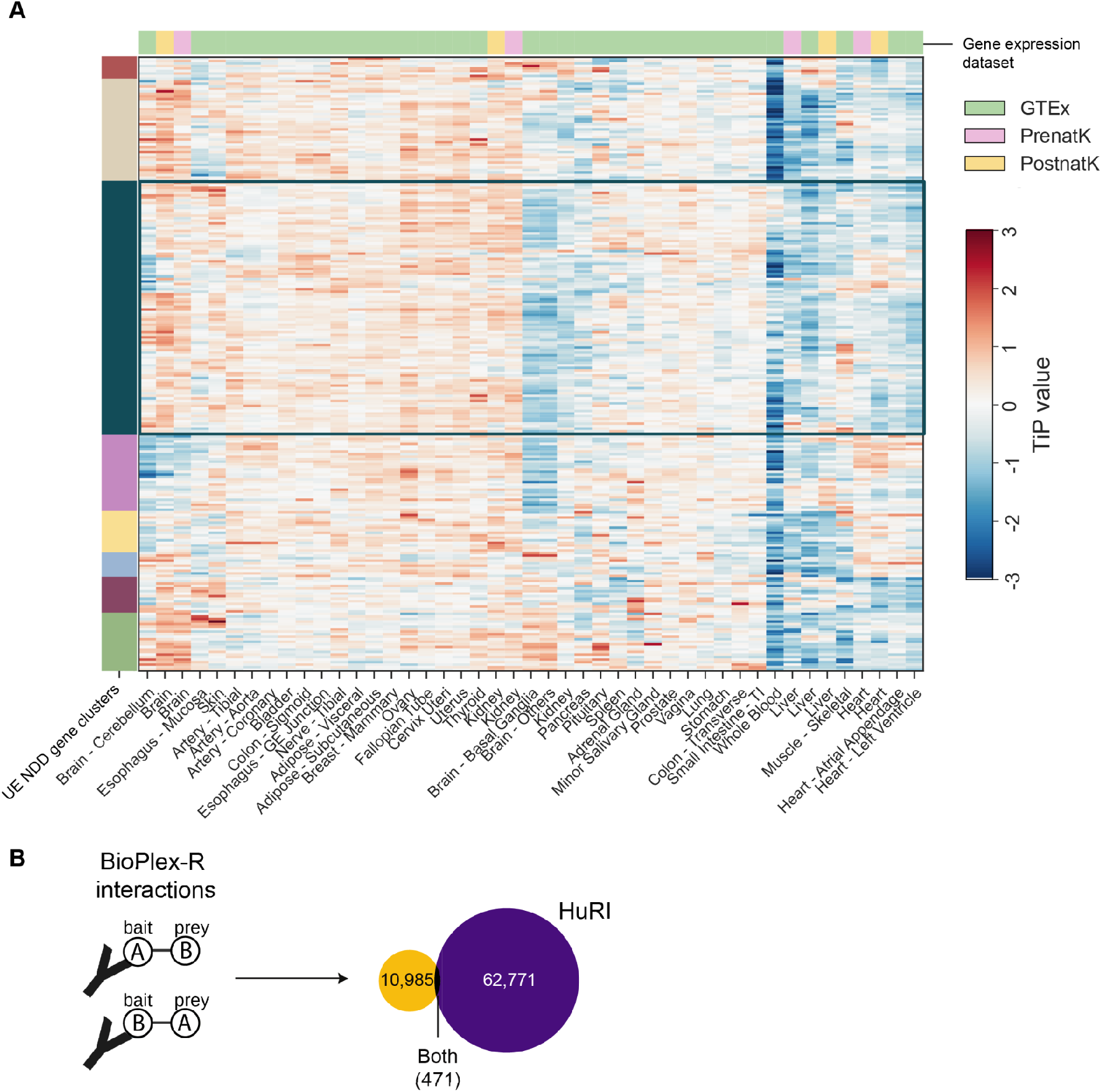
Characterization of uniformly expressed NDD genes. **A.** Heatmap of TiP values of genes from the UE cluster from Figure 2A clustered again based on GTEx and Kaessmann data. The cluster of genes selected for further PPI analysis is indicated. **B.** Schematic of interactions that are part of the reciprocal BioPlex dataset (BioPlex-R) and Venn diagram illustrating the overlap of BioPlexR (yellow) with HuRI (purple) in terms of PPIs.

